# Aged Tendons Exhibit Altered Mechanisms of Strain-Dependent Extracellular Matrix Remodeling

**DOI:** 10.1101/2024.01.26.577397

**Authors:** Anthony N. Aggouras, Emma J. Stowe, Samuel J. Mlawer, Brianne K. Connizzo

## Abstract

Aging is a primary risk factor for degenerative tendon injuries, yet the etiology and progression of this degeneration is poorly understood. While aged tendons have innate cellular differences that support a reduced ability to maintain mechanical tissue homeostasis, the response of aged tendons to altered levels of mechanical loading has not yet been studied. To address this question, we subjected young and aged murine flexor tendon explants to various levels of *in vitro* tensile strain. We first compared the effect of static and cyclic strain on matrix remodeling in young tendons, finding that cyclic strain is optimal for studying remodeling *in vitro*. We then investigated the remodeling response of young and aged tendon explants after 7 days of varied mechanical stimulus (stress-deprivation, 1%, 3%, 5%, or 7% cyclic strain) via assessment of tissue composition, biosynthetic capacity, and degradation profiles. We hypothesized that aged tendons would show muted adaptive responses to changes in tensile strain and exhibit a shifted mechanical setpoint, at which the remodeling balance is optimal. Interestingly, we found 1% cyclic strain best maintains native physiology while promoting ECM turnover for both age groups. However, aged tendons display fewer strain-dependent changes, suggesting a reduced ability to adapt to altered levels of mechanical loading. This work has significant impact in understanding the regulation of tissue homeostasis in aged tendons, which can inform clinical rehabilitation strategies for treating elderly patients.

## INTRODUCTION

Tendons are constantly under loads imposed by muscular contractions during movement, and this stimulus is in turn essential for tendon health [1,2]. Mechanical loads on the whole tissue are transmitted through the hierarchical extracellular matrix (ECM) structure to the cells that reside within it. Resident tendon cells, tenocytes, then sense alterations in their mechanical environment through cell-cell and cell-ECM interactions and can alter their structure and composition to meet functional needs of the tissue. This dynamic, feedback-driven process that maintains tissue homeostasis requires a delicate balance of matrix degradation and synthesis. Tenocytes produce matrix-degrading enzymes, such as matrix metalloproteinases (MMPs), to clear damaged or unneeded matrix proteins. In parallel, tenocytes synthesize new proteins that are incorporated into the matrix and re-organized into functional tissue structure. It has previously been established that moderate mechanical loading, such as exercise, improves tissue function through increases in matrix synthesis [3–8]. Chronic overloading, on the other hand, can shift the balance to catabolic processes, characterized by high breakdown and increased inflammation [9–14]. Therefore, the appropriate regulation of this homeostatic balance of ECM turnover is critical to preventing injury and chronic disease.

Tendon explants enable us to assess extracellular matrix remodeling within an isolated tissue while preserving cell-cell and cell-ECM interactions. As explants can be harvested from donors with different ages, sexes, and genetic backgrounds, explant models can address differences between tissue structure and cell behavior independently. Additionally, mechanical loading of tendon explants with bioreactor systems allows for precise control of the mechanical environment experienced by tissues over the culture period. However, defining the appropriate mechanical stimulation to mimic physiological loading and maintain tissue homeostasis *ex vivo* has proven to be a challenge over the past decade. Multiple studies have worked towards establishing optimal loading protocols to maintain the tendon *ex vivo* and establish a mechanical setpoint that simulates physiological, as well as pathological (ex. overload, fatigue) loading conditions [5,12,15]. However, findings appear to be highly dependent on biological variables, such as tendon site or species, and experimental variables, such as loading rates and modes. Regardless, early work has established that stress-deprivation (SD), complete mechanical unloading, causes a degenerative response, while low level mechanical loading can better maintain native phenotypes [8,12,15,16].

It’s been hypothesized that the process of aging itself could lead to a loss in tissue homeostasis. Aging is a primary risk factor for degenerative tendon injuries; however, the progression and biological drivers of age-related tendon degeneration are poorly understood. Despite many studies on the alterations in mechanical properties, tissue composition, and matrix organization of aged tendons, there is no consensus on specific age-related changes, with significant results often depending on the tendon type and exact donor age of tissue samples [17]. However, it is clear that aged tendons exhibit alterations in cell-mediated processes that support a compromised ability of aged cells to regulate tissue homeostasis [18]. Specifically, aged tendons show decreases in cell density and cellular activity (proliferation, metabolism, matrix synthesis) [16,17,19]. Additionally, multiple studies have documented impaired healing in aged tendons, suggesting an altered ECM repair capacity [20–22]. There has also been reported dysregulation of cell-cell communication in aged tendon stem cells, suggesting a reduced ability to elicit a coordinated tissue-wide remodeling response [23]. Despite this extensive work, the effect of aging on the ability of cells to sense and respond to changing mechanical loads is still largely unexplored. In a previous study from our group, we investigated the response of young and aged tendon explants to stress-deprivation [16]. Despite no age-related differences at baseline, we found that aged tendons show an altered response to the mechanical unloading injury with reduced metabolic activity, proliferation, and matrix biosynthesis. However, the response of aged tendons to altered mechanical demands and the ability to adapt accordingly has not yet been studied.

The objective of this study was to (1) identify optimal loading protocols to stimulate physiological ECM turnover *ex vivo* in both young and aged tendon explants, and (2) investigate the effect of aging on strain-dependent mechanisms of extracellular matrix remodeling. We hypothesized that aged tendon explants would display a muted adaptive response to changes in tensile strain and exhibit a shifted mechanical setpoint from that of young tendons.

## METHODS

### Sample Preparation

Flexor digitorum longus (FDL) tendon explants were harvested from the limbs of young (4 months) and aged (22 months) C57BL/6J male mice directly following sacrifice per approved animal use protocol (BU IACUC PROTO202000046). Following previously described methods [14], all explants were washed in 1x PBS supplemented with 100 units/mL penicillin G, 100 µg/mL streptomycin (Fisher Scientific, Waltham, MA), and 0.25 µg/mL Amphotericin B (Sigma-Aldrich). All explants were then immediately loaded into grips with a 10-mm gauge length (with the intrasynovial segment of the FDL situated between the grips) and placed into a custom-built tensile loading bioreactor (Figure 1A). Throughout culture, explants were kept in culture medium consisting of low glucose Dulbecco’s Modified Eagle’s Media (1 g/l; Fisher Scientific) supplemented with 10% fetal bovine serum (Cytiva, Marlborough, MA), 100 units/mL penicillin G, 100 µg/mL streptomycin (Fisher Scientific), and 0.25 µg/mL Amphotericin B (Sigma-Aldrich). Medium was changed every 2 days in culture for up to 7 days.

**Figure 1.**
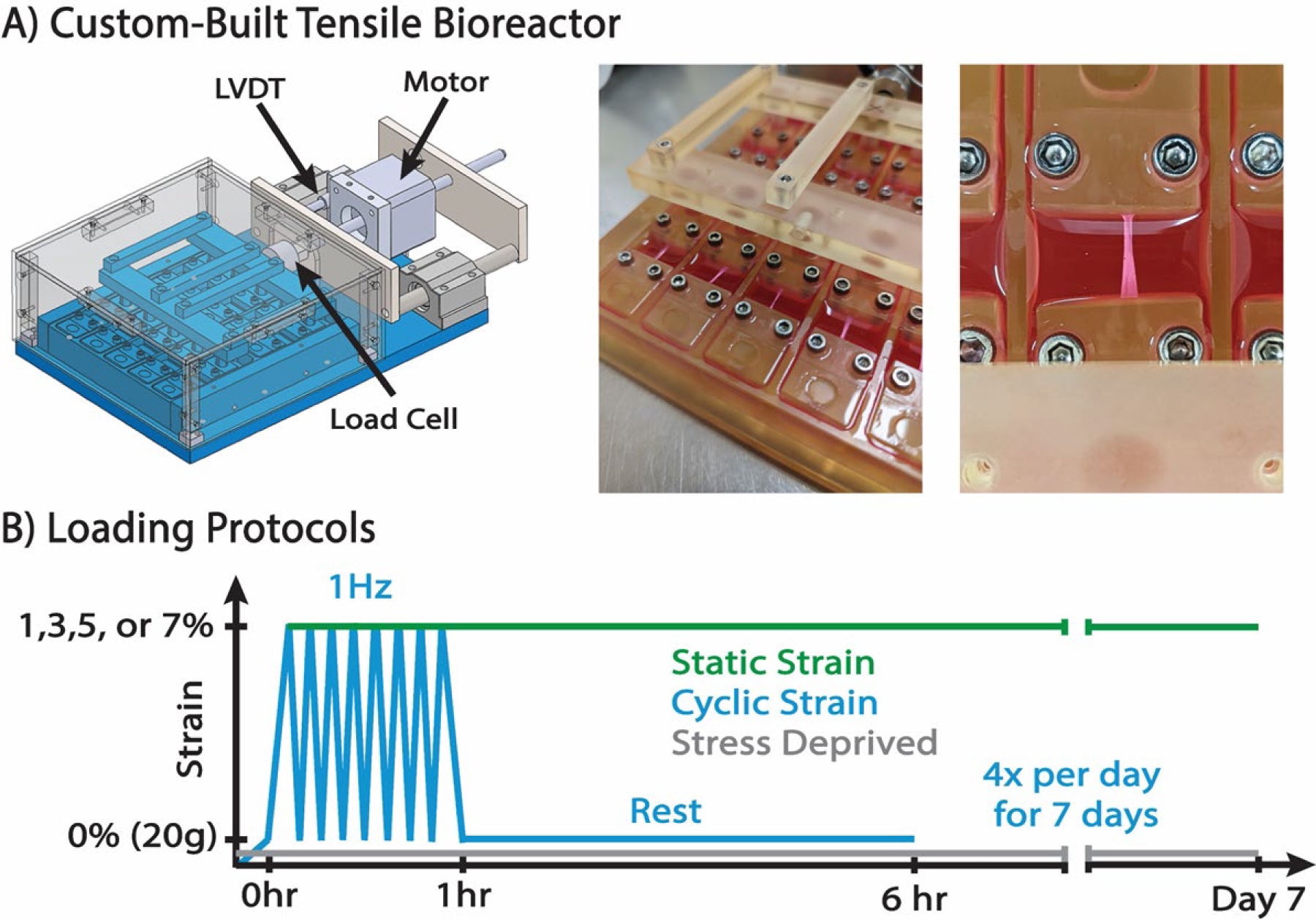
(A) Custom-designed bioreactor (left) and experimental setup of gripped tendon explants (right). (B) Static strain tendons were loaded on day 0 and held at 3, 5%, or 7% maximum strain for the duration of the culture. Cyclic strain (CS) groups were loaded using a triangle waveform from 0% to either 1, 3, 5, or 7% strain at 1 Hz for 1 hour followed by a 5 hour hold at 0% strain. This protocol was repeated 4x a day for 7-days. Stress deprived explants were gripped and left slack for the culture period.

### In Vitro Mechanical Stimulation

Our custom-designed bioreactor system allows for direct control over the mechanical environment experienced by tendon explants. This incubator-housed, tensile-loading bioreactor includes a load cell to measure force and an LVDT to record displacements in real time (Figure 1A). We first explored the effect of loading mode (static vs. cyclic strain) on tendon composition and ECM turnover in young tendon explants only. Then, we investigated the response of both young and aged tendon explants subjected to various levels of cyclic tensile strain. Tendon explants were preloaded to 20g to ensure consistent loading between all explants; this was set as 0% strain. Static strain (SS) tendons were loaded on day 0 and held at their respective strain level for the duration of the culture. Cyclic strain (CS) groups were loaded using a triangle waveform from 0% to either 1, 3, 5, or 7% strain at 1 Hz for 1 hour followed by a 5 hour hold at 0% strain. This protocol was repeated 4 times a day for the duration of the culture (Figure 1B). All tendons were loaded within 1 hour of harvest. An additional group was subjected to stress deprivation (SD), where the explants were gripped but held slack for the duration of the culture period.

### Metabolism, Biosynthesis, and Composition

Explant cell metabolism was measured using a resazurin reduction assay, as previously described [24]. Following a 3-hour incubation with resazurin (diluted 1:10 in culture medium), intensity of the reduced product, resorufin, was measured in collected medium using excitation/emission of 554/584 nm. Values were normalized to daily control wells without explants, such that a value of 1 is representative of a culture without a viable explant. Synthesis of sulfated glycosaminoglycans (sGAG) and total protein (indicative of collagen synthesis) was measured by 24 hour incorporation of ^35^S-sulfate (20 Ci/ml) and ^3^H-proline (10 Ci/ml), respectively (Perkin-Elmer, Norwalk, CT). After culture, explant (n= 5-10/group) wet weight was determined by soaking the tendons in 1x PBS for 1 minute, dabbing excess PBS on a paper towel, and taking the weight of the tendon. This weighing procedure was done in triplicate and the average value of the three weights was considered as the wet weight. The tendons were then lyophilized for at least three hours and dry weights were taken in triplicate. Water content of the tendon was calculated as the difference in wet weight and dry weight divided by the dry weight and multiplied by 100. Following weights, explants were digested overnight with proteinase K (5 mg/mL) (Sigma-Aldrich, St. Louis, MO). Radiolabel incorporation was measured in tissue digests using a liquid scintillation counter (Perkin-Elmer) and incorporation rate was determined. sGAG content was measured using the dimethyl methylene blue (DMMB) assay [25]. Double stranded DNA content was measured using the PicoGreen dye binding assay [26]. A 100 µL portion of each explant digest was then hydrolyzed using 12 M HCl, dried, resuspended, and assayed to measure total collagen content using the hydroxyproline (OHP) assay [27].

### MMP Activity

Activity of matrix metalloproteinases (MMPs) (1,2,3,7,8,9,10,13,14) was determined via analysis of spent culture medium (n= 8-10/group) using a commercially available FRET-based generic MMP cleavage kit (SensoLyte 520 Generic MMP Activity Kit Fluorimetric, Anaspec, Fremont, CA). MMP activity is represented as the concentration of MMP cleaved product (5-FAM-Pro-Leu-OH), the final product of the MMP enzymatic reaction.

### Quantitative Gene Expression

Explants were harvested from each group at day 0 (baseline) and day 7 (n=5-6/group/day). Explants were immediately flash frozen with liquid nitrogen and stored at −80°C until RNA extraction. Samples were placed in Trizol reagent, homogenized with a bead homogenizer (Benchmark Scientific), and then separated using phase-gel tubes (Qiagen)[28]. The supernatant was then purified according to the Zymo Quick-RNA purification kit protocol (Zymo Research). The RNA was then converted into cDNA with reverse transcription and qPCR was performed with the Applied Biosystems StepOne Plus RT-PCR (Applied Biosystems, Foster City, CA). Primer pairs and sequences are listed in the Supplemental Data (Supplemental Table S1). We measured genes responsible for matrix synthesis (*Col1a1*), regulation of collagen fibrillogenesis (*Fmod, Dcn, Bgn*), matrix degradation (*Mmp3, Mmp8, Mmp9, Mmp13*), as well as markers of injury (*Il6, Il1b, Tnfa, Casp3*). Expression for each gene was calculated from the threshold cycle (Ct) value and was normalized to the housekeeping gene β-Actin. All data is represented in log space.

### Data and Statistics

All data are presented as individual data points with summary statistics of mean ± 95% confidence interval. Biosynthesis and composition data are normalized to tendon dry weight to account for any size difference between samples. Data points more than 2 standard deviations outside of the mean were removed as outliers. Statistical evaluation on this set of data was performed using one-way ANOVAs. Bonferroni corrected post-hoc t-tests were then used to identify differences from day 0 metrics and differences from the stress deprived group. For all comparisons, significance was noted at *p<0.05.

## RESULTS

The influence of strain mode (static vs. cyclic) on matrix turnover of young tendons was assessed after 7 days of culture. Explant metabolic activity was increased for both strain modes regardless of the strain level (Figure 2A). At 5% CS, metabolic activity is significantly higher than SS. CS at all levels increased MMP activity whereas static loading maintained low MMP levels (Figure 2B). Protein synthesis was induced by all strain levels across both strain modes (Figure 3C). Regardless of strain level, cyclic loading induced a more robust increase in collagen content compared to SS, which better maintained baseline levels of collagen content (Figure 2D). sGAG synthesis was also induced by all strain levels across both strain modes (Figure 2E), and SS induced greater sGAG incorporation than CS at the 5% strain level only. GAG content was decreased from day 0 levels with CS regardless of strain level, whereas SS maintained GAG content through the 7 days of culture (Figure 2F). While we did not see any notable differences, we did assess gene expression changes between static and cyclic strain modes after 7 days in culture (Supplemental Figure S3).

**Figure 2.**
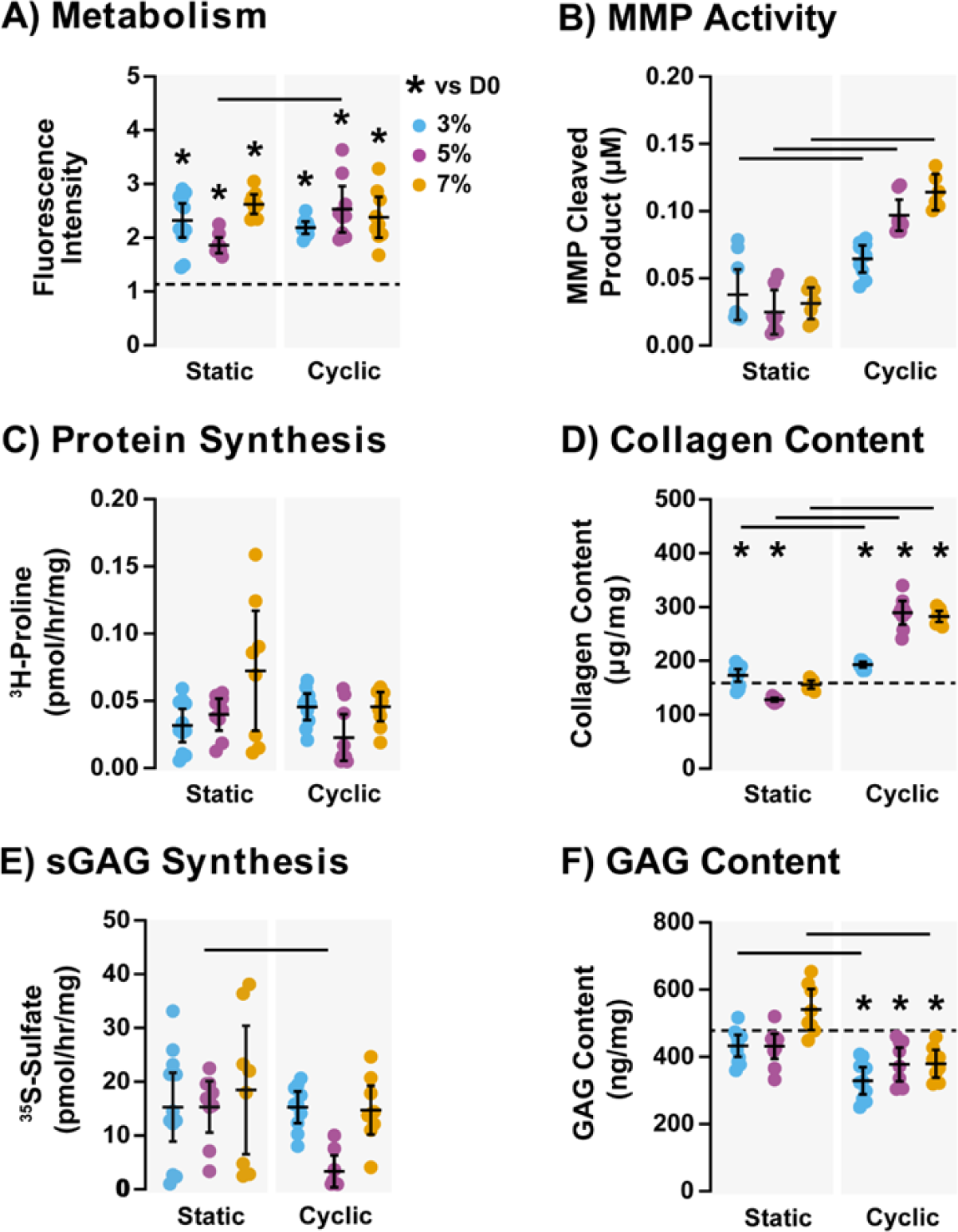
(A) Metabolic activity, (B) MMP activity, (C) total protein synthesis, (D) collagen content, (E) sGAG synthesis, and (F) GAG content of young explants maintained with static strain (left) and cyclic strain (right) after 7 days in culture under 3%, 5%, or 7% strain conditions. Data is presented as individual data points with mean ± 95% confidence interval. Bar (-) indicates significant difference between strain modes at given strain level (p<0.05). Asterisk (*) indicates significant differences from day 0 baseline data (p<0.05), which is represented on graphs by a dotted line.

**Figure 3.**
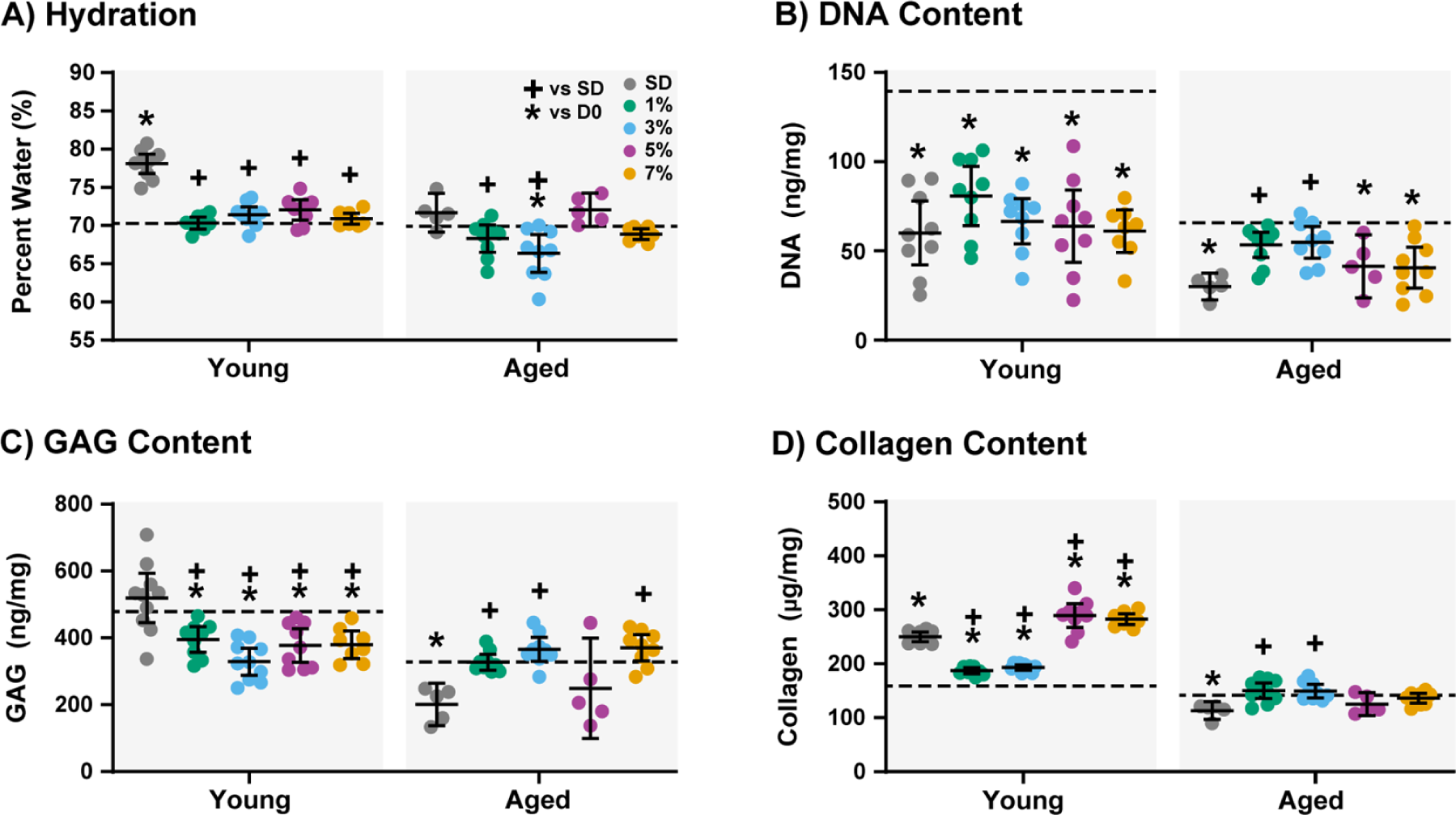
(A) Hydration, (B) DNA content, (C) GAG content, and (D) collagen content of young (left) and aged (right) explants after 7 days in culture under stress-deprivation (SD), 1% cyclic strain, 3% cyclic strain, 5% cyclic strain, or 7% cyclic strain conditions. Data is presented as individual data points with mean ± 95% confidence interval. Plus sign (+) indicates significant difference from SD (p<0.05). Asterisk (*) indicates significant differences from day 0 baseline data (p<0.05), which is represented on graphs by a dotted line.

We next moved to investigate the differences in the response of young and aged tendon explants to various levels of cyclic tensile strain. At baseline, directly following tissue harvest, young FDL explants have more DNA content (indicative of cellularity), GAG content, and collagen content than aged explants (Supplemental Figure S1). There is also greater expression of *Col1a1* and *Fmod* and reduced expression of *Dcn*, *Mmp3*, *Il1b*, and *Tnfa* at baseline in aged tissues (Supplemental Figure S2).

The ECM remodeling response of young and aged explants was then assessed after a week under cyclic loading only. Water content is maintained by all strain levels in the young tendons, and this is significantly different from stress deprivation where tissues are more hydrated after culture (Figure 3A). In aged samples, all loading protocols maintain baseline values except for 3% strain. In the young explants, all loading protocols result in a decrease of DNA content over culture (Figure 3B). However, in the aged explants, the DNA content is maintained with both the 1 and 3% groups (Figure 3B). Cyclic loading of young tendons resulted in a decrease in GAG content over culture, whereas aged explants maintained GAG content (Figure 3C). For collagen content, all loading protocols result in an increase in collagen content in young tendons and a maintenance of collagen content in aged tendons (Figure 3D).

Metabolic activity was increased from baseline for every group regardless of age (Figure 4A). In the young explants, the increase in metabolic activity increases with strain, peaking at 5% CS. However, in the aged explants, 1% CS showed the highest metabolic activity. Protein synthesis, which is indicative of collagen synthesis, is more responsive to changes in CS loading conditions in young explants (Figure 4B). The young explants have lower matrix protein synthesis in the 1% and 5% CS groups when compared to the SD group. The aged explants are all at a low level of protein synthesis regardless of the magnitude of strain applied to the tendons during culture. For sGAG synthesis, all tendons but 5% had moderate levels of sGAG synthesis (Figure 4C). For the aged tendons, sGAG synthesis appears to increase with increasing strain level and is increased compared to SD at 3% and 7% CS. MMP activity, which is indicative of matrix degradation, increases with greater strain magnitudes in young tendons (Figure 4D). In the aged tendons, high strains do not initiate high MMP activity, as seen in young samples.

**Figure 4.**
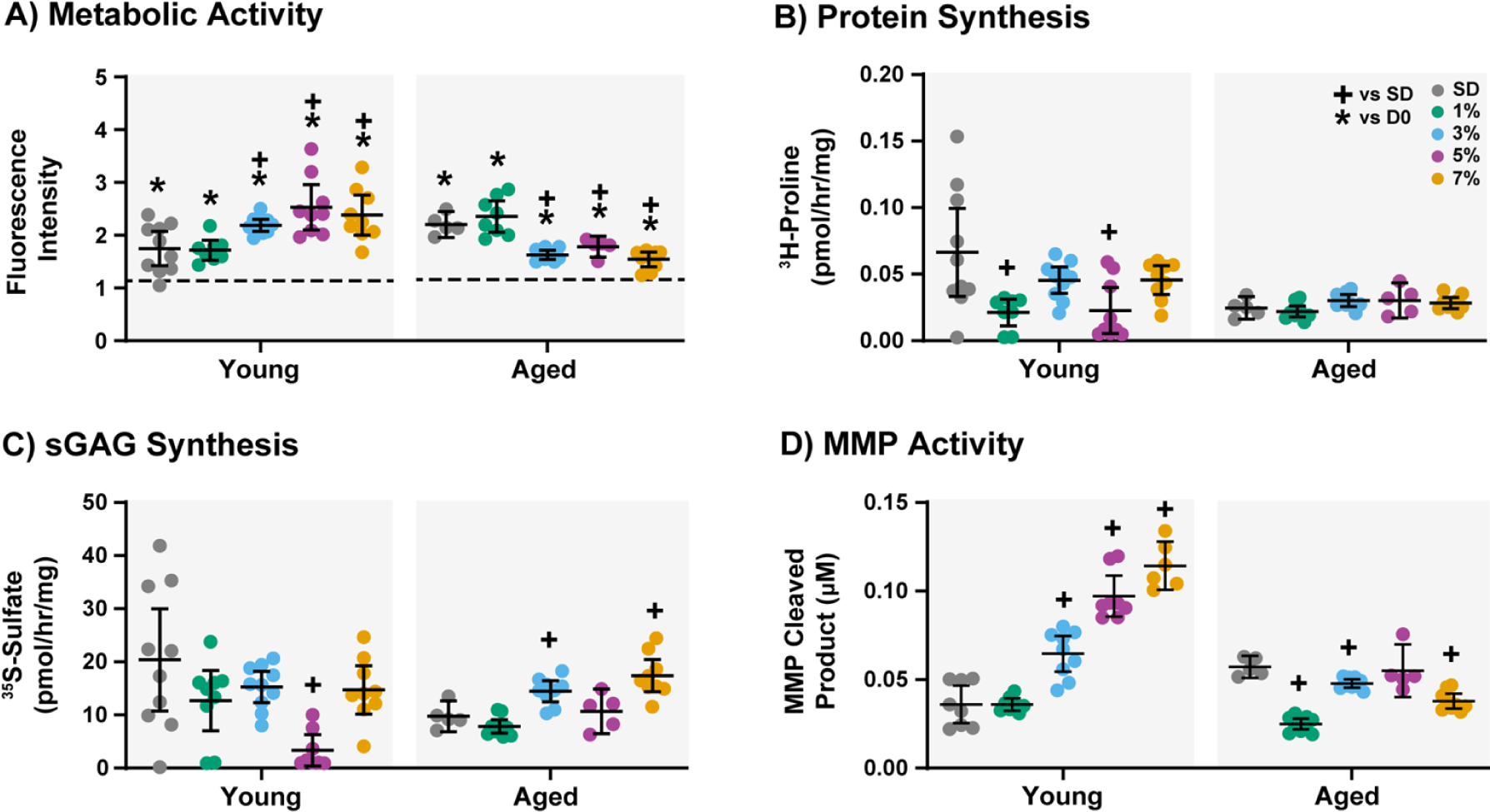
(A) Metabolic activity, (B) total protein synthesis, (C) sGAG synthesis, and (D) MMP activity of young (left) and aged (right) explants after 7 days in culture under stress-deprivation (SD), 1% cyclic strain, 3% cyclic strain, 5% cyclic strain, or 7% cyclic strain conditions. Data is presented as individual data points with mean ± 95% confidence interval. Plus sign (+) indicates significant difference from SD (p<0.05). Asterisk (*) indicates significant differences from day 0 baseline data (p<0.05), which is represented on graphs by a dotted line.

We then looked at changes in expression of genes associated with matrix turnover after 7 days of culture. In the young explants, baseline *Col1a1* expression is only maintained by the 7% cyclic group (Figure 5A). *Col1a1* expression is increased in the young SD, 3% CS, and 5% CS groups and decreased in the young 1% CS group. In the aged explants, *Col1a1* expression is downregulated compared to the baseline day 0 expression at every strain level. For *Fmod* expression, young explants show downregulation from baseline for every strain level except 5% CS (Figure 5B). In aged explants, all strain levels except 3% CS are downregulated compared to baseline. Young explants maintain day 0 *Dcn* expression at 1, 3, and 5% CS levels but 7% CS caused a decrease in expression (Figure 5C). In aged tendons, baseline *Dcn* expression is maintained over the 7 days of culture regardless of strain level. For *Bgn*, the young explants maintain day 0 expression over the 7 days of culture regardless of CS level (Figure 5D). However, in the aged tendons, *Bgn* expression is downregulated with SD, 1%, and 7 % CS levels and maintained with 3 and 5% CS.

**Figure 5.**
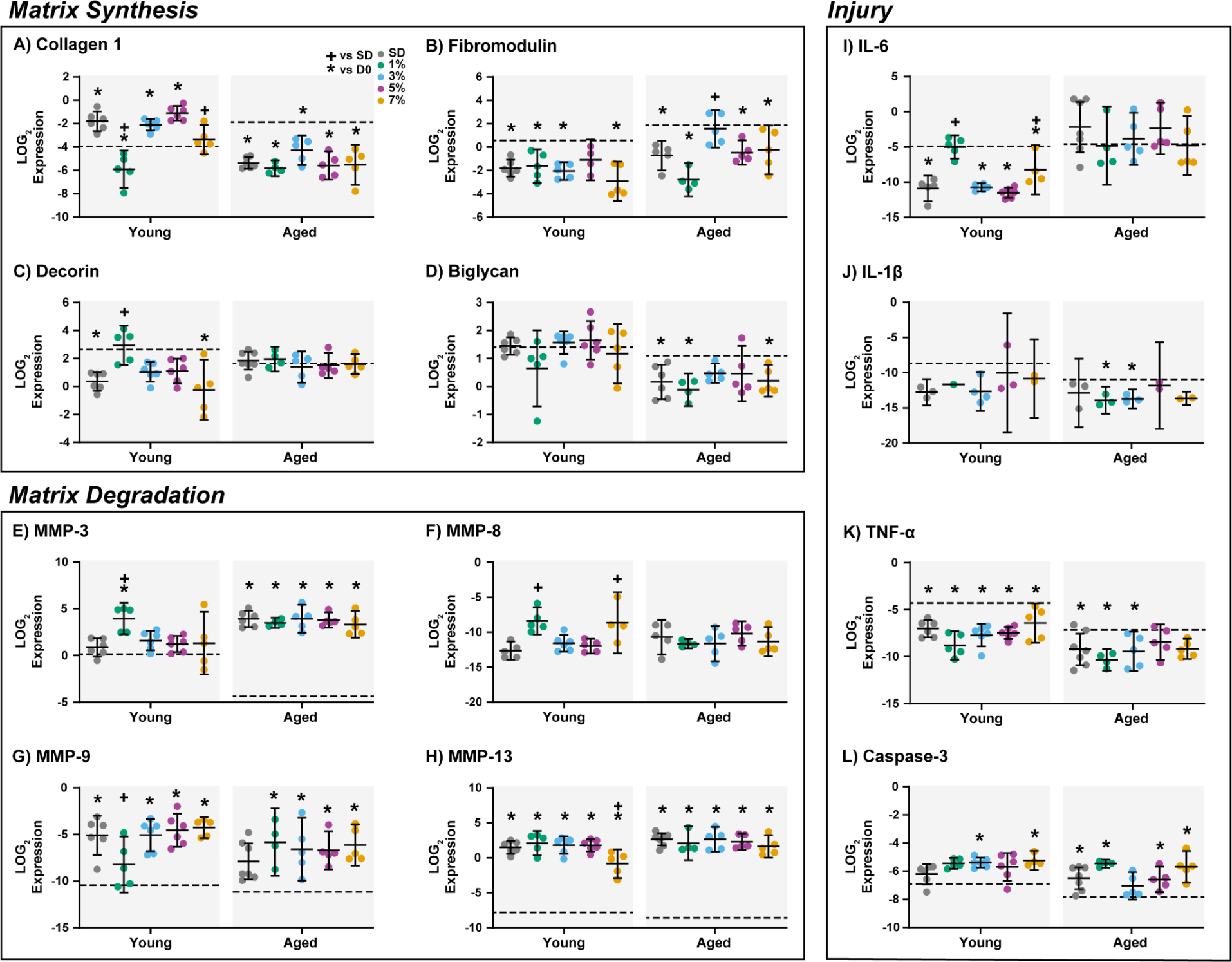
Gene expression of (A-D) extracellular matrix proteins, (E-H) matrix metalloproteinases, and (I-L) injury markers of young (left) and aged (right) explants after 7 days in culture under stress-deprivation (SD), 1% cyclic strain, 3% cyclic strain, 5% cyclic strain, or 7% cyclic strain conditions. Data is presented as individual data points with mean ± 95% confidence interval. Plus sign (+) indicates significant difference from SD (p<0.05). Asterisk (*) indicates significant differences from day 0 baseline data (p<0.05), which is represented on graphs by a dotted line.

Next we considered genes that are involved in regulating matrix degradation. In young explants, *Mmp3* expression was only upregulated relative to baseline for the 1% CS strain, whereas in the aged tendons, all strain levels upregulated *Mmp*3 expression substantially (Figure 5E). For *Mmp8*, there were no detectable transcript levels at day 0 for either young or aged explants. While there are no strain-dependent differences in *Mmp8* expression in young tendons at day 7, there is increased *Mmp8* expression in 1% and 7% CS (Figure 5F). For *Mmp9*, all groups are significantly upregulated from the baseline expression except in the 1% CS group for the young explants and the SD group for the aged explants (Figure 5G). For *Mmp13*, all strain levels induce a significant increase in expression after 7 days of culture (Figure 5H).

Finally, we evaluated the expression of genes that serve as injury markers after 7 days of culture. In young explants, *Il6* expression was maintained at day 0 expression only in the 1% CS group; all other strain levels had a significant downregulation in *Il6* expression (Figure 5I). In aged explants, baseline *Il6* expression was maintained regardless of the strain level. For *Il1b*, young explants maintained day 0 expression regardless of the strain level (Figure 5J). In aged tendons, *Il1b* expression was downregulated in the 1 and 3% CS groups. For *Tnfa* expression, young explants exhibited a downregulation regardless of strain level (Figure 5K). Aged explants exhibited a downregulation at SD, 1%, 3% CS levels. *Casp3* expression, indicative of apoptosis, was upregulated in young explants subjected to 3% and 5% CS compared to baseline (Figure 5L). In aged explants, only the 3% strain group is able to maintain day 0 expression levels and all other strain levels exhibit an increase in Casp3 expression.

## DISCUSSION

Tendon explants are a powerful model system that enable direct interrogation of matrix remodeling processes *ex vivo*; however, a major challenge of tendon explant culture has been identifying the precise loading conditions that promote homeostasis, matrix remodeling, or matrix damage (injury) [29]. Mechanical loading state depends on a number of factors including frequency, magnitude, loading duration, rest period, and loading modality. We sought to determine which strain mode was the best at stimulating murine FDL tendons to adapt to a range of physiological strain levels. Typically, tendons *in vivo* are loaded in the linear range of their stress-strain curve with physiological strains ranging from 2-6% [30,31]. Previous research has indicated that in tendon and tendon fascicle explants, physiological cyclic loading between 4-6% strain at 0.25-1 Hz leads to tendon remodeling with increased collagen production during culture [11,12,32]. While these studies have indicated that cyclic loading conditions may be best for inducing matrix remodeling in tendon explants, they have been conducted in larger animal models and in different tendons, including Achilles tendons and rat tail tendon fascicles. The extensibility of tendons in response to physiological loads is both tendon- and species-dependent [33]. Therefore, the strain modes and strain levels that promote matrix remodeling in the mouse flexor tendon may be different then previously studied models. This knowledge gap compelled us to establish whether static or cyclic strain better supports tendon matrix remodeling during explant culture.

Our data suggests that cyclic loading, regardless of strain level, induces turnover and matrix remodeling. We can see that both metabolic activity and matrix synthesis are induced across strain levels (3-7%) and both strain modes. Seven days of explant culture is sufficient to induce tenocyte mediated synthesis of fibrillar collagen matrix, indicated by ^3^H-proline incorporation, and the synthesis of non-fibrillar matrix proteins such as proteoglycans, indicated by ^35^S-sulfate incorporation. While matrix synthesis is comparable between the strain modes, matrix degradation, the other key mechanism of matrix turnover, is more sensitive to cyclic strain. Whereas static loading maintained low MMP levels, cyclic strain at all levels increased activity of MMPs, which are important to the tenocytes ability to cleave and remove damaged or unneeded matrix [34]. It’s possible that static strain does not result in as much microdamage formation as cyclic strain, and therefore increased MMP activity is less necessary. Finally, CS has a larger effect on explant tissue composition. Regardless of strain level, cyclic loading induced collagen incorporation and GAG loss whereas static loading maintained collagen and GAG content. This would suggest that while static strain is able to support synthesis of matrix proteins and the production of MMPs, it is unable to induce compositional change to the tendon explant over the 7 days of culture. With lower levels of matrix turnover throughout culture, static strain explants may have been more quiescent synthetically, resulting in maintenance of matrix composition. We also have to keep in mind that this current study is only out to 7 days of culture and it is possible that static strain wouldn’t be ideal for long term studies. Furthermore, long duration at high strains may start to become injurious to the tendon. For studying how ECM turnover mechanisms are altered with conditions such as exercise, injury, or aging, cyclic loading protocols appear to be more optimal, as they promote higher levels of tissue remodeling and are more physiologically similar to *in vivo* loading.

One of the primary goals of this project was to establish age-specific mechanical setpoints by identifying optimal loading conditions that stimulate *in vitro* matrix remodeling. We assessed this by identifying which cyclic strain magnitude best maintained the baseline tendon physiology for each of the biomarkers assessed in this study. These conclusions are summarized in Figure 6, where we established the mechanical setpoint of the tendons to be the lowest magnitude of cyclic strain that most closely maintains the baseline physiology of the tendon for each age group. We found that 1% cyclic strain best maintains tendon explants *in vitro,* regardless of age. Previous work has primarily explored cyclic strain protocols in young tendon explants [5,12,15,32,35,36]. These studies lack a consensus on the optimal strain to maintain tendons *in vitro*, primarily because each study is conducted in a different tendon model with inconsistencies in number of loading cycles, duration of loading, and length of culture. Despite the variability in methodology, together these studies indicate that a low level of strain (1-6%) is sufficient to preserve the tendon during culture. This diversity in strain level is likely due to the variety of anatomical roles of the studied tendons, which can have substantial influence on the biomechanical response [33]. For positional tendons, such as the FDL, as examined here, the optimal strain was identified to be around 1% and within the range of the toe region of the stress-strain curve [15,29,35].

**Figure 6.**
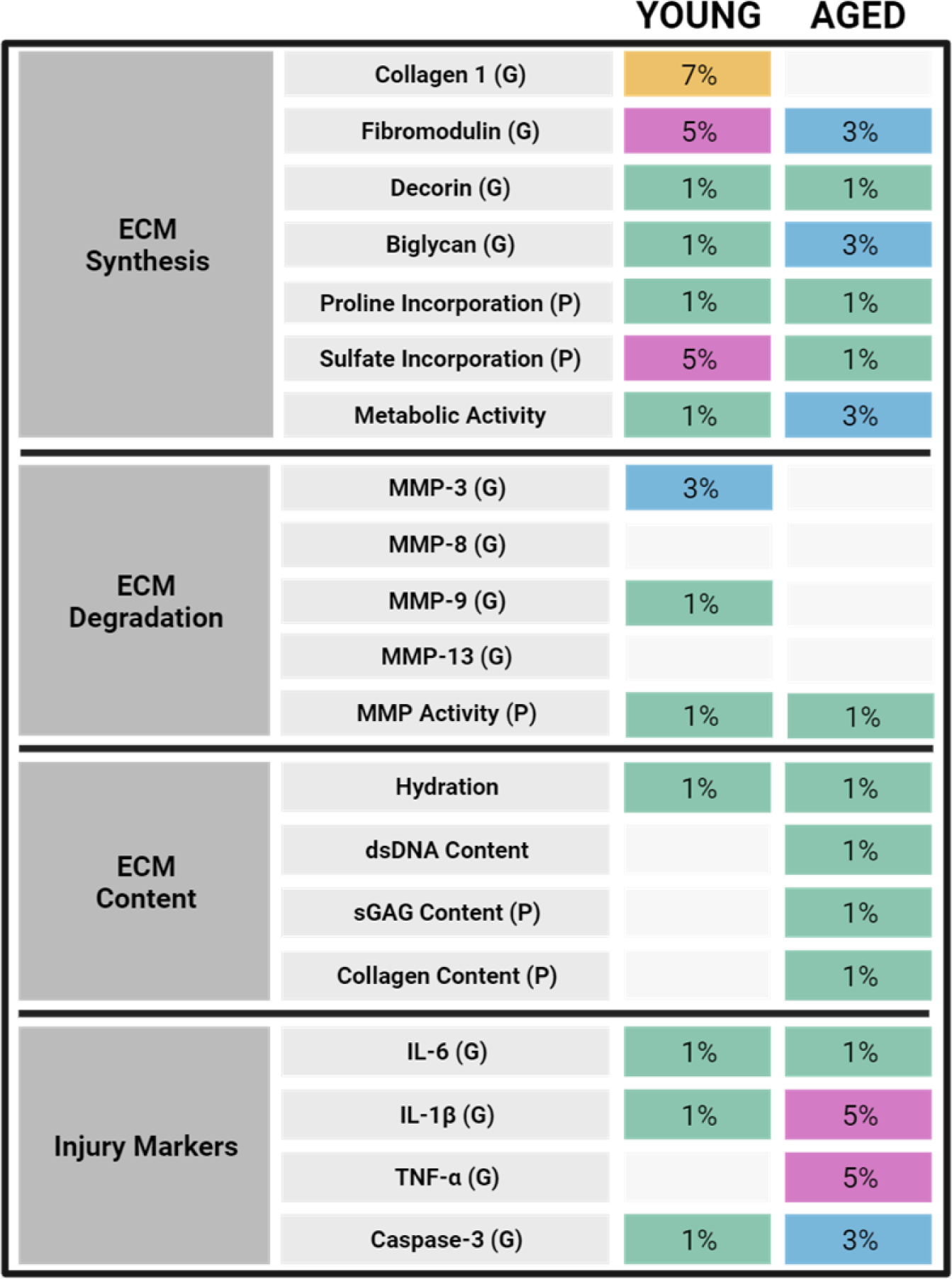
Summary figure indicating the lowest magnitude of cyclic strain that most closely maintains baseline physiology. Blank boxes signify that none of the tested cyclic loading protocols could maintain native conditions. (G) indicates gene expression and (P) indicates protein level biomarker.

While our conclusions about 1% cyclic strain as an optimal loading protocol for tendon explants is aligned with previous work, it is important to note some additional considerations with the mechanical loading protocols used in this set of studies. We recognize that load-controlled mechanical loading has been suggested to be more physiological relevant [37]; however, we chose to utilize a stain-controlled bioreactor system because of the technical simplicity of the system and better control of mechanical parameters. Using our strain-controlled protocols we have demonstrated that a low level of cyclic strain is sufficient to maintain tendon physiology while also inducing physiological mechanisms of matrix turnover. Despite this we believe there is room for further optimization of our cyclic loading protocol, including further assessment of rest periods and cycle numbers. In the future, we hope to confirm similar findings using load-controlled mechanical actuation.

While young and aged tendons exhibit no differences in their homeostatic mechanical setpoints, we do observe that aged tenocytes exhibit desensitized remodeling responses to altered levels of cyclic strain compared to young tenocytes. Examining tissue composition after 7-days of culture with optimal strain conditions, we see significant deviations in content (DNA, GAG, collagen) from baseline levels in young tissues, but not in aged tissues. In young tissues, we document increased collagen content and decreased GAG content over the culture period, indicating both a sensitivity to cyclic mechanical loading and an initiation of protective remodeling processes. The failure of aged tissues to elicit a similar response suggests a loss of adaptive mechanisms that allows the tissue to meet changing mechanical demands, thus increasing the risk of injury. We also document an interesting trend in strain-dependent metabolic activity in young and aged tissues. In young tissues, metabolic activity increases with increasing strain, signifying an increased metabolic demand to adapt to higher mechanical loads. Aged tissues do not experience comparable increases in metabolic activity, suggesting a failure to produce necessary levels of energy for tissue adaptation.

We also document a significant increase in the activity of degradative enzymes at higher strains in young tissues, which is not replicated in aged tissues. It’s important to note that our generic MMP activity captures all MMP families, not just those specific to collagen breakdown, and that our data only captures late-stage biosynthetic activity (day 7). Therefore, it is possible that differences were present earlier in culture. Future studies should investigate early timepoints to fully determine the time course of strain-dependent degradation and synthesis. Additionally, strain-dependent trends were not consistent between gene and protein assays. Across all strain groups, the expression of *Mmp8, Mmp9,* and *Mmp13* is significantly increased from baseline for both age groups. However, for *Mmp3* which is known to degrade proteoglycans [38], an upregulation from baseline is only seen in aged tissues, suggesting a potential ramp up in proteoglycan degradation. As we see substantial loss of GAG over culture for young but not aged tissues, this potentially indicates a delayed response to GAG breakdown in aged tissues.

Surprisingly, we did not see an upregulation of the injury markers *Il6, Il1b, Tnfa,* or *Casp3* in either the young or aged tendons at any strain level despite the 7% strain being greater than the assumed physiologic range. The lack of injury response may be due to substantial recovery during the rest period or that the tested strains did not reach high enough magnitudes, as previous studies have indicated that cyclic strains above 9% are sufficient to induce injury to the tendon, with matrix and cell damage as well as increased apoptosis [11–14]. It is also possible that the expression of the injury markers at the higher strain levels may have occurred earlier in culture and the expression reduced as culture time progressed. Future studies will aim to identify the stain levels that may be injurious to both young and aged tendon explants by exploring higher strain levels, assessing timepoints earlier in culture, and quantifying cell death and apoptosis.

Regardless, across the multiple metrics of ECM turnover assessed in this work, we document pronounced strain-dependency of young tendon ECM remodeling. The lack of comparable responses in aged tissues signifies a loss of strain-sensitivity and adaptive remodeling with aging. Interestingly, our findings about lack of tensile strain adaptation in aged tissues are similar to what was found recently in another study from our group that examined the response of young and aged tendons to an acute compressive injury [39]. In both studies, while young tissues display significant remodeling changes in response to the various mechanisms of mechanical loading, aged tendons exhibit a more muted response, with little or no remodeling changes. It is possible that this lack of adaptation could be attributed to decreases in cellularity in aged tissues or altered cellular communication [16,21,23]. Previous *in vivo* work has documented mechanosensitive mechanisms in aged human achilles tendons, documenting increased stiffness following 14-weeks of cyclic loading exercise [40]. While we do observe mechanosensitive, strain-dependent changes in our aged tendons, the differences are muted compared to those in young tissues.

Of course, this study is not without its limitations. We recognize that analysis of tendon composition, synthetic activity, and gene expression at day 7 of culture only captures the late stage adaptation of the tendon to altered mechanical conditions. While assessing late stage markers is sufficient to identify the establishment of homeostasis, this analysis lacks the ability to fully characterize distinct stages of cellular responses as well as the mechanisms responsible for age-dependent remodeling outcomes. We also recognize that it is typically common to evaluate tendon function via mechanical assessment [15,32,40] after culture. Unfortunately, we did not have enough samples to perform mechanical testing due to the large number of experiments necessary for the other assays examining specific cell-mediated processes. In the future, we will perform these endpoint assessments as well as utilize real-time load and displacement data collected from our bioreactor system to assess how tendon mechanics change throughout the culture period. To focus the scope of this study we only assessed one loading duration and frequency. While we were able to identify a sufficient strain level to maintain tendon physiology using the protocol described in this study, there could be more optimal loading parameters that would better meet our desired outcomes. Future studies will aim to refine our loading protocol further to optimize tendon health and matrix remodeling during culture.

Despite these limitations, we have utilized *in vitro* tensile loading of murine FDL tendon explants to establish a homeostatic loading protocol of 1% cyclic strain. This gives us the ability to explore pathological conditions of mechanical loading and resulting remodeling profiles in both young and aged tissues. Furthermore, we document a reduced response in aged tendons and a lack of specific strain-dependent adaptations that could contribute to high prevalence of tendon injuries in elderly populations. Future work will continue to explore the ability of young and aged tendons to sense mechanical loads and establish new states of tissue homeostasis, specifically through analysis of mechanical and biological properties before and after step changes in tissue strain. We also hope to tease out individual mechanisms resulting in altered ECM remodeling in aged tissues through investigation of age-related biological processes, such as cellular senescence [41]. Finally, we are now working to evaluate how age-associated remodeling is altered with biological sex and sex hormone levels by performing similar studies in female animals [16].

In light of our findings, this study furthers our understanding of the regulation of tissue homeostasis in aged and young tendons during mechanical loading. Importantly, this work is the first to confirm that young and aged tendons display altered mechanisms of matrix remodeling in response to tensile loading, supporting the notion that tendon pathologies require age-specific clinical interventions and therapies. Furthermore, this study establishes a strong model for future work to explore mechanosensitive matrix remodeling mechanisms that promote matrix adaptation or pathological degeneration. These explorations into age-specific cellular responses to *in vitro* mechanical loading will provide fundamental insights into the regulation of tissue homeostasis in aged tendons, which can inform clinical rehabilitation strategies for treating elderly patients.

## ACKNOWLEDGMENT

We would like to thank Elliot Frank for his expertise with the tensile bioreactor system and Henry Chow for his assistance in explant culture experiments.

## NOMENCLATURE

Anova: Analysis of Variance
*Bgn*: biglycan
*Casp3*: Caspase 3
*Col1a1*: collagen, type I, alpha 1
*CS*: Cyclic Strain
*Dcn*: decorin
ECM: Extracellular Matrix
FDL: Flexor Digitorum Longus
*Fmod*: fibromodulin
GAG: Glycosaminoglycan
*Il1b*: interleukin 1 beta
*Il6*: interleukin 6
MMP: Matrix Metalloproteinase
*Mmp3*: matrix metallopeptidase 3
*Mmp8*: matrix metallopeptidase 8
*Mmp9*: matrix metallopeptidase 9
*Mmp13*: matrix metallopeptidase 13
PBS: Phosphate Buffered Saline
*SD*: Stress Deprivation
sGAG: Sulfated Glycosaminoglycan
*SS*: Static Strain
*Tnfa*: tumor necrosis factor alpha

---

**Supplemental Figure 1:**
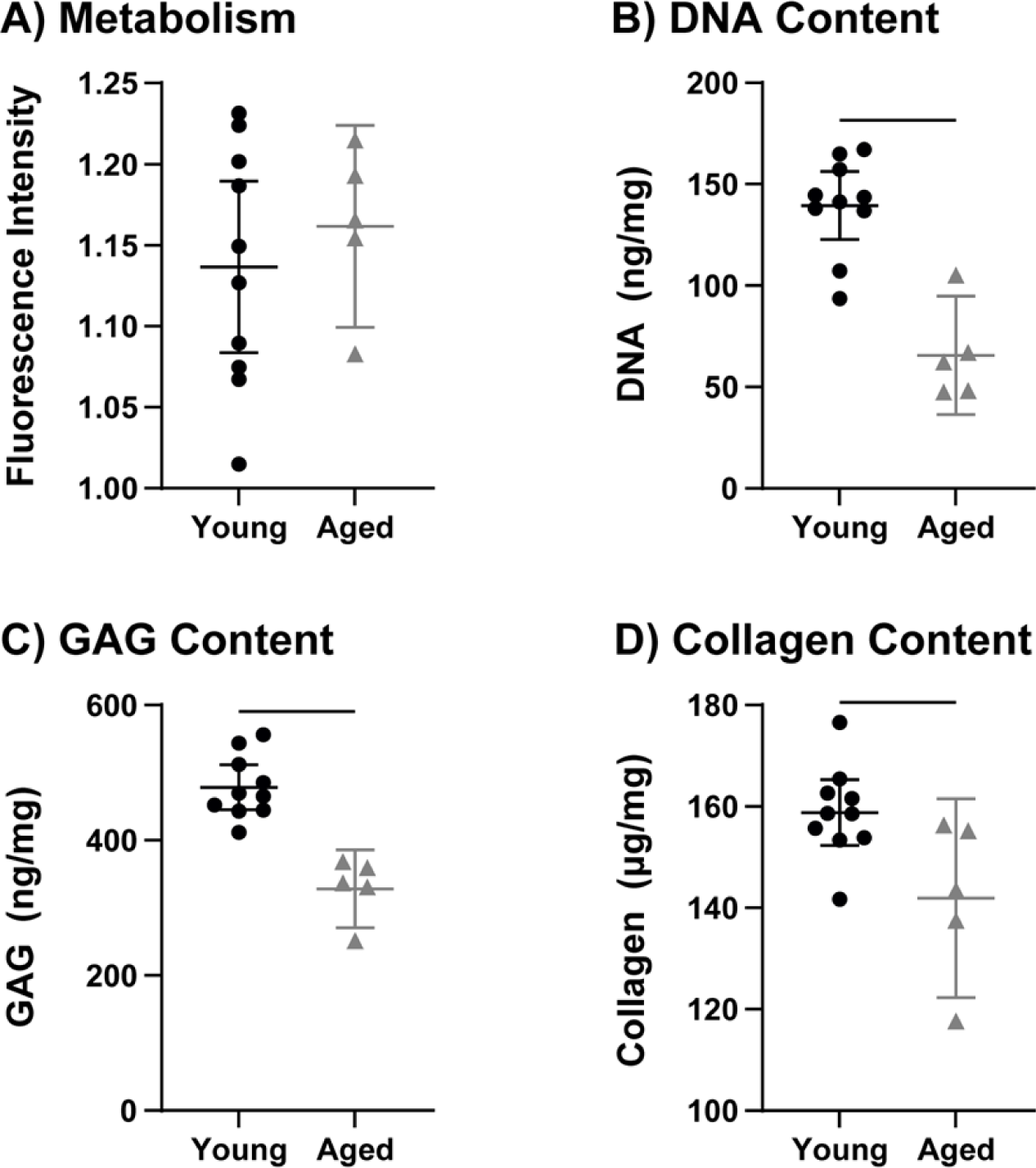
(A) Metabolism, (B) DNA content, (C) GAG content, and (D) collagen content at baseline (day 0) in young and aged flexor explants. Data is presented as individual data points with mean ± 95% confidence interval. Bar (-) spanning between groups indicates significant difference (p<0.05).

**Supplemental Figure 2:**
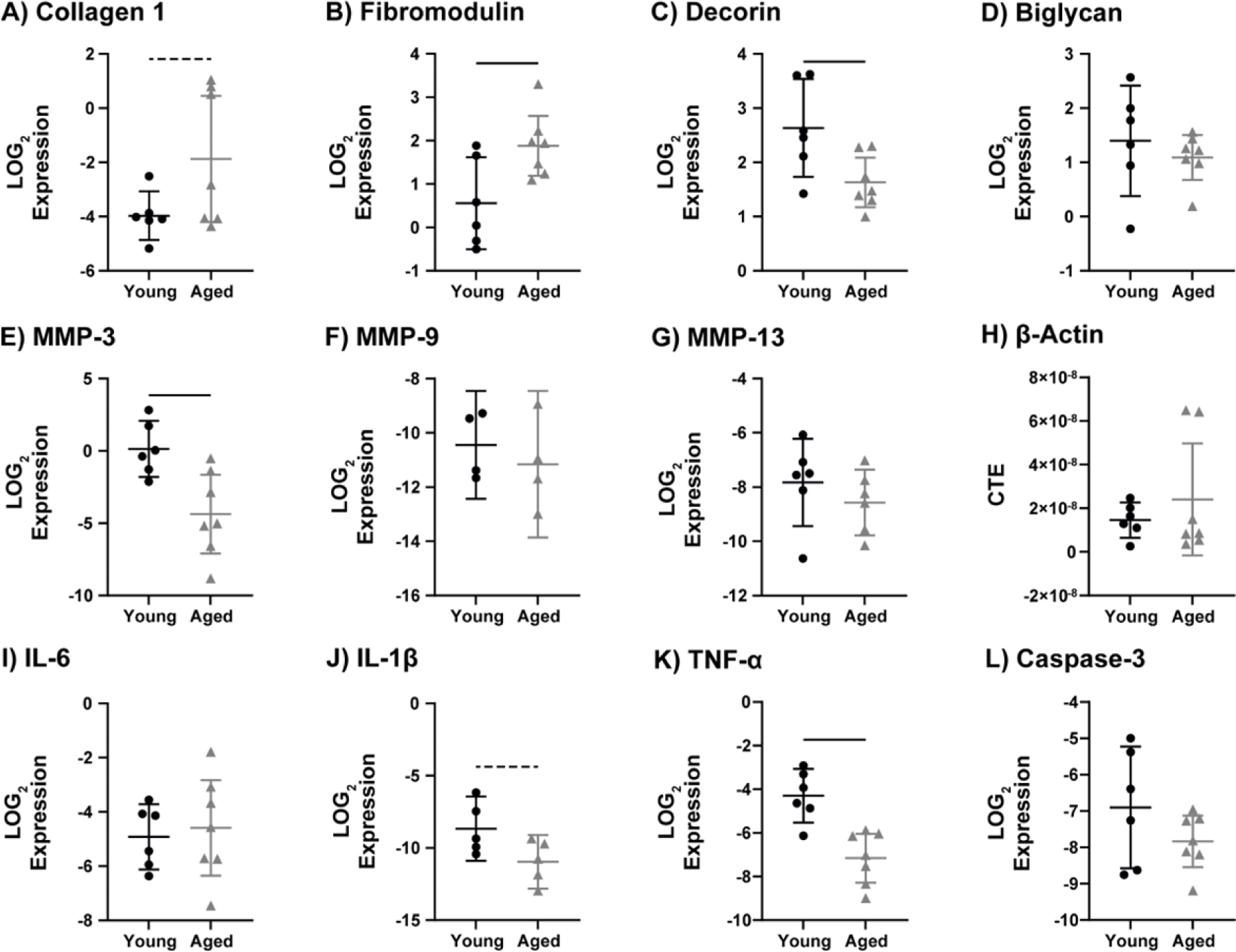
Gene expression of (A) *Col1a1*, (B) *Fmod*, (C) *Dcn*, (D) *Bgn*, (E) *Mmp3*, (F) *Mmp9*, (G) *Mmp13*, (H) *Bact*, (I) *Il6*, (J) *Il1b*, (K) *Tnfa*, and (L) *Casp*3 at baseline (day 0) in young and aged flexor explants. Data is presented as individual data points with mean ± 95% confidence interval. Bar (-) spanning between groups indicates significant difference (p<0.05).

**Supplemental Figure 3:**
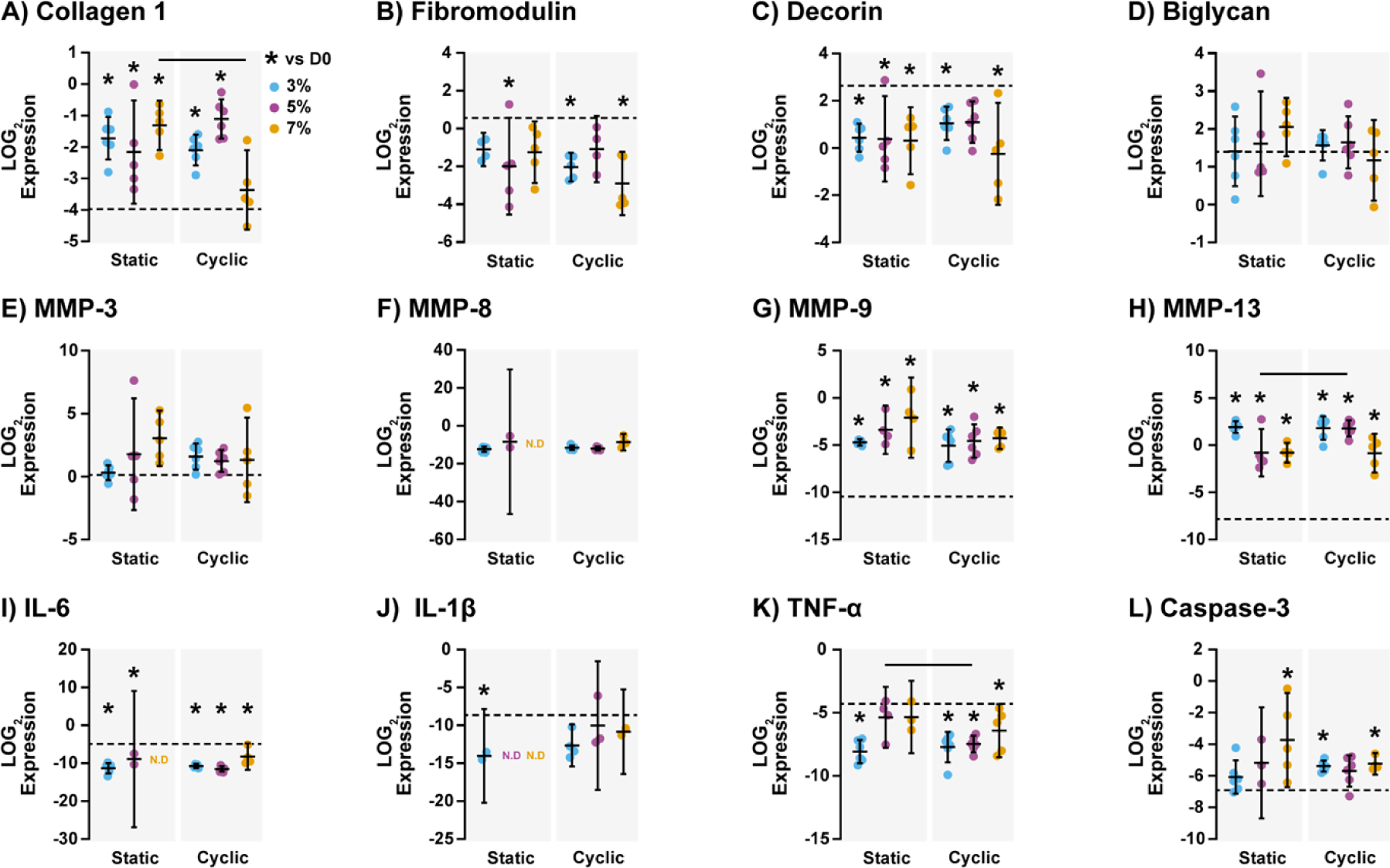
Gene expression of (a-d) extracellular matrix proteins, (e-f) matrix metalloproteinases, and (i-l) injury markers of young explants maintained with static strain (left) and cyclic strain (right) after 7 days in culture under 3%, 5%, or 7% strain conditions. Data is presented as individual data points with mean ± 95% confidence interval. Bar (-) indicates significant difference between strain modes at given strain level (p<0.05). Asterisk (*) indicates significant differences from day 0 baseline data (p<0.05), which is represented on graphs by a dotted line.

**Supplemental Table S1.**
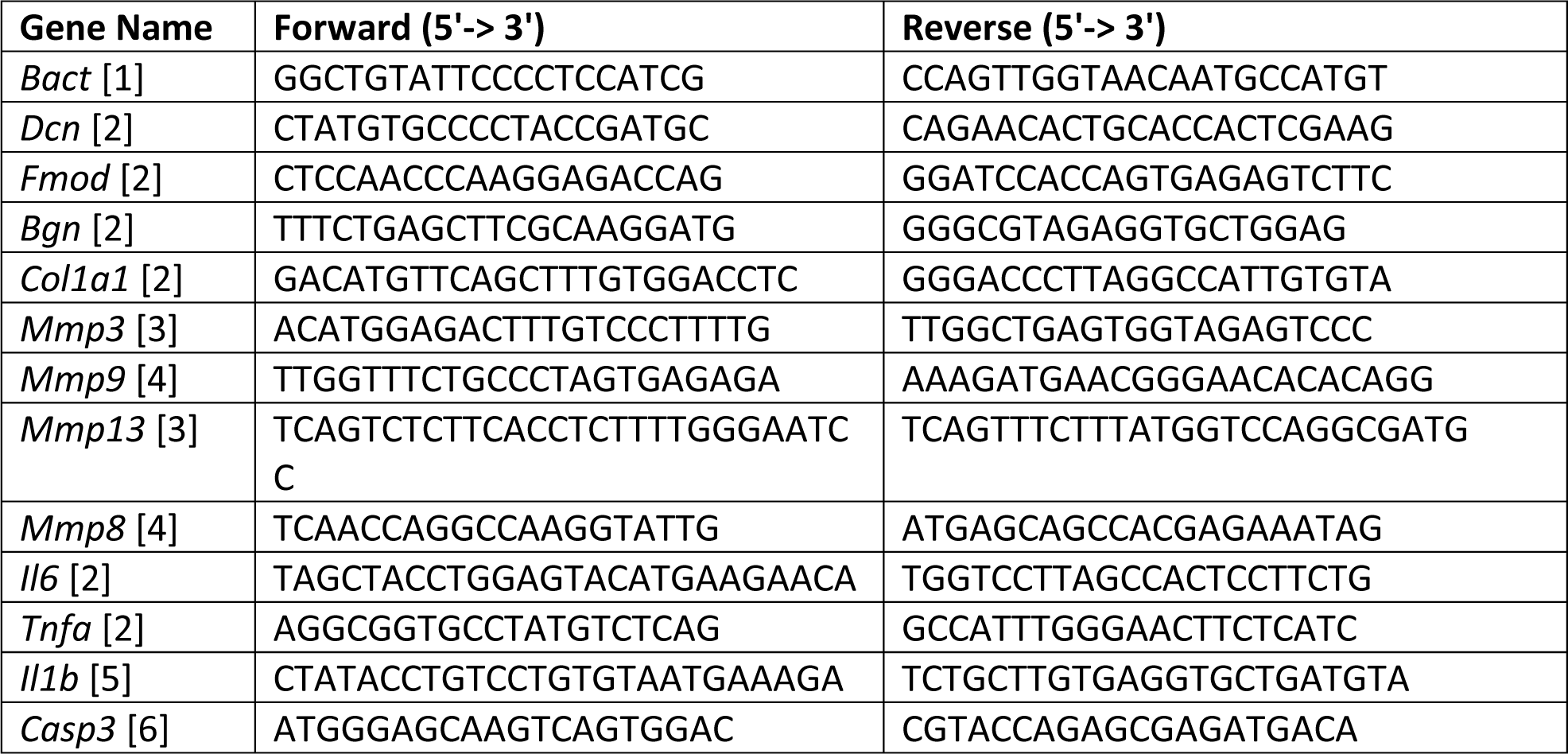
PCR primer forward and reverse sequences used for quantitative gene expression analysis.

## REFERENCES

[1] Maganaris, C. N., and Paul, J. P., 1999, “In Vivo Human Tendon Mechanical Properties,” The Journal of Physiology, 521(1), pp. 307–313.

[2] Bojsen-Møller, J., and Magnusson, S. P., 2015, “Heterogeneous Loading of the Human Achilles Tendon In Vivo,” Exercise and Sport Sciences Reviews, 43(4), p. 190.

[3] Bohm, S., Mersmann, F., and Arampatzis, A., 2015, “Human Tendon Adaptation in Response to Mechanical Loading: A Systematic Review and Meta-Analysis of Exercise Intervention Studies on Healthy Adults,” Sports Medicine - Open, 1(1).

[4] Rooney, S. I., Tobias, J. W., Bhatt, P. R., Kuntz, A. F., and Soslowsky, L. J., 2015, “Genetic Response of Rat Supraspinatus Tendon and Muscle to Exercise,” PLOS ONE, 10(10), p. e0139880.

[5] Screen, H. R. C., Shelton, J. C., Bader, D. L., and Lee, D. A., 2005, “Cyclic Tensile Strain Upregulates Collagen Synthesis in Isolated Tendon Fascicles,” Biochemical and Biophysical Research Communications, 336(2), pp. 424–429.

[6] Heinemeier, K. M., Olesen, J. L., Haddad, F., Langberg, H., Kjaer, M., Baldwin, K. M., and Schjerling, P., 2007, “Expression of Collagen and Related Growth Factors in Rat Tendon and Skeletal Muscle in Response to Specific Contraction Types,” The Journal of Physiology, 582(3), pp. 1303–1316.

[7] Abreu, E. L., Leigh, D., and Derwin, K. A., 2008, “Effect of Altered Mechanical Load Conditions on the Structure and Function of Cultured Tendon Fascicles,” J Orthop Res, 26(3), pp. 364–73.

[8] Arnoczky, S. P., Tian, T., Lavagnino, M., and Gardner, K., 2004, “Ex Vivo Static Tensile Loading Inhibits MMP-1 Expression in Rat Tail Tendon Cells through a Cytoskeletally Based Mechanotransduction Mechanism,” J. Orthop. Res., 22(2), pp. 328–333.

[9] Spiesz, E. M., Thorpe, C. T., Chaudhry, S., Riley, G. P., Birch, H. L., Clegg, P. D., and Screen, H. R., 2015, “Tendon Extracellular Matrix Damage, Degradation and Inflammation in Response to in Vitro Overload Exercise,” Journal of Orthopaedic Research, 33(6), pp. 889–97.

[10] Fung, D. T., Wang, V. M., Andarawis-Puri, N., Basta-Pljakic, J., Li, Y., Laudier, D. M., Sun, H. B., Jepsen, K. J., Schaffler, M. B., and Flatow, E. L., 2010, “Early Response to Tendon Fatigue Damage Accumulation in a Novel in Vivo Model,” J Biomech, 43(2), pp. 274–279.

[11] Legerlotz, K., Jones, G. C., Screen, H. R., and Riley, G. P., 2013, “Cyclic Loading of Tendon Fascicles Using a Novel Fatigue Loading System Increases Interleukin-6 Expression by Tenocytes,” Scand J Med Sci Sports, 23(1), pp. 31–7.

[12] Wang, T., Lin, Z., Day, R. E., Gardiner, B., Landao-Bassonga, E., Rubenson, J., Kirk, T. B., Smith, D. W., Lloyd, D. G., Hardisty, G., Wang, A., Zheng, Q., and Zheng, M. H., 2013, “Programmable Mechanical Stimulation Influences Tendon Homeostasis in a Bioreactor System,” Biotechnol. Bioeng., 110(5), pp. 1495–1507.

[13] Szczesny, S. E., Aeppli, C., David, A., and Mauck, R. L., 2018, “Fatigue Loading of Tendon Results in Collagen Kinking and Denaturation but Does Not Change Local Tissue Mechanics,” J Biomech, 71, pp. 251–256.

[14] Scott, A., Khan, K. M., Heer, J., Cook, J. L., Lian, O., and Duronio, V., 2005, “High Strain Mechanical Loading Rapidly Induces Tendon Apoptosis: An Ex Vivo Rat Tibialis Anterior Model,” Br J Sports Med, 39(5), p. e25.

[15] Wunderli, S. L., Widmer, J., Amrein, N., Foolen, J., Silvan, U., Leupin, O., and Snedeker, J. G., 2018, “Minimal Mechanical Load and Tissue Culture Conditions Preserve Native Cell Phenotype and Morphology in Tendon—a Novel Ex Vivo Mouse Explant Model,” Journal of Orthopaedic Research, 36(5), pp. 1383–1390.

[16] Connizzo, B. K., Piet, J. M., Shefelbine, S. J., and Grodzinsky, A. J., 2020, “Age-Associated Changes in the Response of Tendon Explants to Stress Deprivation Is Sex-Dependent,” Connect Tissue Res, 61(1), pp. 48–62.

[17] Siadat, S. M., Zamboulis, D. E., Thorpe, C. T., Ruberti, J. W., and Connizzo, B. K., 2021, “Tendon Extracellular Matrix Assembly, Maintenance and Dysregulation Throughout Life,” Adv Exp Med Biol, 1348, pp. 45–103.

[18] Korcari, A., Przybelski, S. J., Gingery, A., and Loiselle, A. E., 2022, “Impact of Aging on Tendon Homeostasis, Tendinopathy Development, and Impaired Healing,” Connective Tissue Research, 0(0), pp. 1–13.

[19] Magnusson, S. P., and Kjaer, M., 2019, “The Impact of Loading, Unloading, Ageing and Injury on the Human Tendon,” J Physiol, 597(5), pp. 1283–1298.

[20] Mienaltowski, M. J., Dunkman, A. A., Buckley, M. R., Beason, D. P., Adams, S. M., Birk, D. E., and Soslowsky, L. J., 2016, “The Injury Response of Geriatric Mouse Patellar Tendons,” J Orthop Res, 34(7), pp. 1256–1263.

[21] Ackerman, J. E., Bah, I., Jonason, J. H., Buckley, M. R., and Loiselle, A. E., 2017, “Aging Does Not Alter Tendon Mechanical Properties during Homeostasis, but Does Impair Flexor Tendon Healing,” J. Orthop. Res., 35(12), pp. 2716–2724.

[22] Lai, F., Tang, H., Wang, J., Lu, K., Bian, X., Wang, Y., Shi, Y., Guo, Y., He, G., Zhou, M., Zhang, X., Zhou, B., Zhang, J., Chen, W., and Tang, K., 2021, “Effects of Aging on the Histology and Biochemistry of Rat Tendon Healing,” BMC Musculoskelet Disord, 22, p. 949.

[23] Popov, C., Kohler, J., and Docheva, D., 2017, “Activation of EphA4 and EphB2 Reverse Signaling Restores the Age-Associated Reduction of Self-Renewal, Migration, and Actin Turnover in Human Tendon Stem/Progenitor Cells,” Front. Aging Neurosci., 7.

[24] Connizzo, B. K., and Grodzinsky, A. J., 2018, “Release of Pro-Inflammatory Cytokines from Muscle and Bone Causes Tenocyte Death in a Novel Rotator Cuff in Vitro Explant Culture Model,” Connect Tissue Res, 59(5), pp. 423–436.

[25] Farndale, R. W., Buttle, D. J., and Barrett, A. J., 1986, “Improved Quantitation and Discrimination of Sulphated Glycosaminoglycans by Use of Dimethylmethylene Blue,” Biochim. Biophys. Acta, 883(2), pp. 173–177.

[26] Singer, V. L., Jones, L. J., Yue, S. T., and Haugland, R. P., 1997, “Characterization of PicoGreen Reagent and Development of a Fluorescence-Based Solution Assay for Double-Stranded DNA Quantitation,” Anal. Biochem., 249(2), pp. 228–238.

[27] Dourte, L. M., Pathmanathan, L., Mienaltowski, M. J., Jawad, A. F., Birk, D. E., and Soslowsky, L. J., 2013, “Mechanical, Compositional, and Structural Properties of the Mouse Patellar Tendon with Changes in Biglycan Gene Expression,” J. Orthop. Res., 31(9), pp. 1430–1437.

[28] Grinstein, M., Dingwall, H. L., Shah, R. R., Capellini, T. D., and Galloway, J. L., 2018, “A Robust Method for RNA Extraction and Purification from a Single Adult Mouse Tendon,” PeerJ, 6, p. e4664.

[29] Wunderli, S. L., Blache, U., and Snedeker, J. G., 2020, “Tendon Explant Models for Physiologically Relevant Invitro Study of Tissue Biology - a Perspective,” Connect Tissue Res, 61(3–4), pp. 262–277.

[30] Wang, T., Chen, P., Zheng, M., Wang, A., Lloyd, D., Leys, T., Zheng, Q., and Zheng, M. H., 2018, “In Vitro Loading Models for Tendon Mechanobiology,” Journal of Orthopaedic Research, 36(2), pp. 566–575.

[31] Benage, L. G., Sweeney, J. D., Giers, M. B., and Balasubramanian, R., 2022, “Dynamic Load Model Systems of Tendon Inflammation and Mechanobiology,” Front Bioeng Biotechnol, 10, p. 896336.

[32] Wang, T., Lin, Z., Ni, M., Thien, C., Day, R. E., Gardiner, B., Rubenson, J., Kirk, T. B., Smith, D. W., Wang, A., Lloyd, D. G., Wang, Y., Zheng, Q., and Zheng, M. H., 2015, “Cyclic Mechanical Stimulation Rescues Achilles Tendon from Degeneration in a Bioreactor System,” Journal of Orthopaedic Research, 33(12), pp. 1888–1896.

[33] Kjær, M., 2004, “Role of Extracellular Matrix in Adaptation of Tendon and Skeletal Muscle to Mechanical Loading,” Physiological Reviews, 84(2), pp. 649–698.

[34] Del Buono, A., Oliva, F., Osti, L., and Maffulli, N., 2013, “Metalloproteases and Tendinopathy,” Muscles Ligaments Tendons J, 3(1), pp. 51–57.

[35] Tohidnezhad, M., Zander, J., Slowik, A., Kubo, Y., Dursun, G., Willenberg, W., Zendedel, A., Kweider, N., Stoffel, M., and Pufe, T., 2020, “Impact of Uniaxial Stretching on Both Gliding and Traction Areas of Tendon Explants in a Novel Bioreactor,” Int J Mol Sci, 21(8).

[36] Maeda, E., Shelton, J. C., Bader, D. L., and Lee, D. A., 2007, “Time Dependence of Cyclic Tensile Strain on Collagen Production in Tendon Fascicles,” Biochemical and Biophysical Research Communications, 362(2), pp. 399–404.

[37] Pedaprolu, K., and Szczesny, S. E., 2022, “A Novel, Open-Source, Low-Cost Bioreactor for Load-Controlled Cyclic Loading of Tendon Explants,” Journal of Biomechanical Engineering, 144(084505).

[38] Cui, N., Hu, M., and Khalil, R. A., 2017, “Biochemical and Biological Attributes of Matrix Metalloproteinases,” Prog Mol Biol Transl Sci, 147, pp. 1–73.

[39] Mlawer, S. J., Frank, E. H., and Connizzo, B. K., “Aged Tendons Lack Adaptive Response to Acute Compressive Injury,” Journal of Orthopaedic Research, n/a(n/a).

[40] Epro, G., Mierau, A., Doerner, J., Luetkens, J. A., Scheef, L., Kukuk, G. M., Boecker, H., Maganaris, C. N., Brüggemann, G.-P., and Karamanidis, K., 2017, “The Achilles Tendon Is Mechanosensitive in Older Adults: Adaptations Following 14 Weeks versus 1.5 Years of Cyclic Strain Exercise,” Journal of Experimental Biology, 220(6), pp. 1008–1018.

[41] Stowe, E., and Connizzo, B., 2023, “Cellular Senescence Suppresses ECM Synthesis in Response to Mechanical Unloading in Tendon Explants.,” 2023 Summer Biomechanics, Bioengineering, and Biotransport Conference., Vail, Colorado.

## References

[1] Veres-Székely, A., Pap, D., Sziksz, E., Jávorszky, E., Rokonay, R., Lippai, R., Tory, K., Fekete, A., Tulassay, T., Szabó, A. J., and Vannay, Á., 2017, “Selective Measurement of α Smooth Muscle Actin: Why β-Actin Can Not Be Used as a Housekeeping Gene When Tissue Fibrosis Occurs,” BMC Mol Biol, 18(1), p. 12.

[2] Connizzo, B. K., Piet, J. M., Shefelbine, S. J., and Grodzinsky, A. J., 2020, “Age-Associated Changes in the Response of Tendon Explants to Stress Deprivation Is Sex-Dependent,” Connect Tissue Res, 61(1), pp. 48–62.

[3] Connizzo, B. K., and Grodzinsky, A. J., 2018, “Release of Pro-Inflammatory Cytokines from Muscle and Bone Causes Tenocyte Death in a Novel Rotator Cuff in Vitro Explant Culture Model,” Connect Tissue Res, 59(5), pp. 423–436.

[4] Mota, R., Parry, T. L., Yates, C. C., Qiang, Z., Eaton, S. C., Mwiza, J. M., Tulasi, D., Schisler, J. C., Patterson, C., Zaglia, T., Sandri, M., and Willis, M. S., 2018, “Increasing Cardiomyocyte Atrogin-1 Reduces Aging-Associated Fibrosis and Regulates Remodeling in Vivo,” Am J Pathol, 188(7), pp. 1676–1692.

[5] Xuan, Y., Gao, Y., Huang, H., Wang, X., Cai, Y., and Luan, Q. X., 2017, “Tanshinone IIA Attenuates Atherosclerosis in Apolipoprotein E Knockout Mice Infected with Porphyromonas Gingivalis,” Inflammation, 40(5), pp. 1631–1642.

[6] Zulliger, R., Lecaudé, S., Eigeldinger-Berthou, S., Wolf-Schnurrbusch, U. E. K., and Enzmann, V., 2011, “Caspase-3-Independent Photoreceptor Degeneration by N-Methyl-N-Nitrosourea (MNU) Induces Morphological and Functional Changes in the Mouse Retina,” Graefes Arch Clin Exp Ophthalmol, 249(6), pp. 859–869.

